# Metatranscriptomic Profiling of Viral Diversity in Aquatic Environments with a Focus on Microalgal Viruses

**DOI:** 10.64898/2026.11.05.764614

**Authors:** Ravi Ojha, Tatiana Demina, Chiara Beltrani, Maryam Abidizadegan, Heli A. M. Mönttinen, Hanna M. Oksanen, Elina Peltomaa, Yuji Tomaru, Tero Ahola, Mohammadreza Sadeghi

## Abstract

Freshwater microeukaryotes contribute to aquatic productivity, yet the RNA viruses associated with these communities remain incompletely characterized compared with their marine counterparts. Here, we used metatranscriptomics to profile viral diversity across five Finnish freshwater sites (one river and four lakes) and one coastal marine sample from Hiroshima Bay in the Seto Inland Sea, Japan. We recovered 399 viral operational taxonomic units (vOTUs), of which 285 (71.4%) represented RNA viruses distributed across five viral phyla and putatively associated with fungal, plant or algal, metazoan and bacterial hosts. DNA viruses formed a smaller component of the dataset, including bacteriophage-like vOTUs with transcripts encoding replication, structural, packaging, and lysis proteins. Two vOTUs, which were detected in the Vantaanjoki river sample, were putatively classified within the family *Marnaviridae*, which comprises RNA viruses that infect protists in marine and freshwater environments. Both vOTUs encoded conserved replication and picornavirus-like capsid domains, and one represented a near-complete dicistronic genome. RNA-dependent RNA polymerase (RdRp)-based phylogeny placed these vOTUs in a strongly supported lineage outside the currently recognized genera within the family *Marnaviridae*, supporting their provisional assignment to the candidate genus “Vandarnavirus” proposed here within *Marnaviridae.* Whole-genome comparisons using the IMG/VR v4 database revealed genetic similarities between the vOTUs representing the candidate “Vandarnavirus” genus and picorna-like virus sequences from freshwater, wetland, saltmarsh, and estuarine metatranscriptomes. These findings expand the known ecological range of marnavirus-like RNA viruses and identify freshwater habitats as under-sampled reservoirs of microalgae-associated viral sequences.

**Importance:** Freshwater ecosystems support primary production and nutrient cycling, but their associated RNA-virus diversity remains incompletely characterized. Here, metatranscriptomics recovered diverse viral sequences from five Finnish freshwater sites and one exploratory marine sample from Hiroshima Bay, Japan. Two marnavirus-like vOTUs were detected only in the Vantaanjoki river sample and formed a distinct RdRp lineage outside currently recognized *Marnaviridae* genera. We provisionally propose the candidate genus “Vandarnavirus”; formal taxonomic placement and host associations require further evaluation. By connecting freshwater viral discovery with genome architecture and phylogeny, this work expands the known ecological range of marnavirus-like viruses and highlights freshwater habitats as under-sampled reservoirs of microalgal viruses.

## Introduction

Aquatic ecosystems are sustained by diverse microbial communities that drive global carbon and nutrient cycling. Among these, microalgae are major primary producers, underpinning aquatic food webs and contributing substantially to global carbon fixation^1^. In both freshwater and coastal marine systems, these photosynthetic organisms support complex assemblages of bacteria, archaea, microeukaryotes, and viruses that together form dynamic microbial consortia, or holobionts^2,3^. The productivity and stability of these communities are shaped not only by microbial interactions but also by viral infection. Viruses are increasingly recognized as powerful ecological agents that regulate microbial populations, influence bloom dynamics, and redirect organic matter through infection-driven lysis^4–6^. By modulating host abundance and metabolic activity, viruses influence nutrient recycling and energy flow across aquatic ecosystems, making them integral components of microbial ecology^7,8^.

Despite their ecological importance, viruses infecting aquatic microeukaryotes remain comparatively understudied. Research on algal viruses has largely focused on large double-stranded DNA (dsDNA) viruses infecting marine phytoplankton, whereas RNA viruses in aquatic systems have received far less attention^5,9^. Advances in high-throughput sequencing reveal that RNA viruses represent a vast and previously underappreciated component of the global virosphere^10,11^. Many aquatic RNA viruses possess positive-sense single-stranded RNA (+ssRNA) genomes and are affiliated with the order *Picornavirales*^2^. Environmental sequencing surveys have uncovered extensive RNA viral diversity, yet most sequences recovered from environmental datasets lack close relatives among characterized viruses^12,13^. This gap highlights a major limitation in our understanding of RNA virus evolution and ecological roles^14,15^.

Among RNA viruses infecting protists, the family *Marnaviridae* represents one of the best-characterized groups. Members of this family are small, non-enveloped viruses with +ssRNA genomes typically ranging from approximately 8.6 to 9.6 knt. Their genomes encode conserved replication-associated domains, including a helicase and an RNA-dependent RNA polymerase (RdRp), together with capsid proteins^16^. Two principal genome architectures occur within the family: mono-partite, with a single large open reading frame (ORF) encoding a polyprotein, or a bipartite (dicistronic), with separate ORFs encoding replication and structural proteins. Several marnaviruses infect marine microalgae and diatoms, suggesting an important role in regulating phytoplankton populations and microbial community turnover^2,12^. However, most characterized members originate from marine environments, leaving a substantial knowledge gap regarding the diversity, host range, and ecological distribution of marnaviruses in freshwater ecosystems^5,12,16^.

Finland, with its extensive network of lakes and rivers, provides an ideal setting for investigating viral diversity in freshwater ecosystems. In parallel, coastal environments in Japan, such as Hiroshima Bay, are highly productive systems with diverse microalgal assemblages and provide an opportunity to include a geographically distant marine environment as an exploratory external comparison.

Here, we used metatranscriptomic sequencing to characterize viral communities associated with microeukaryotes across Finnish freshwater systems, which included one river and four lake sites, and a coastal marine microalgal habitat within the Seto Inland Sea (Hiroshima Bay, Japan). Because surface-water samples were concentrated using a 20-µm plankton net, this study primarily characterizes viruses recovered from the plankton-retained fraction of the sampled waters rather than the complete aquatic or microalgal virosphere. Viruses associated with pico- and nanoplankton and free viral particles passing through the mesh may therefore be underrepresented. In this study, we aimed to identify and classify RNA viruses recovered from the plankton-retained fraction of these ecosystems and to examine the genome organization and distribution of microalgal-associated RNA viruses belonging to the family *Marnaviridae*.

By reconstructing viral genomes and analyzing their phylogenetic relationships, this study expands current knowledge of aquatic RNA virus diversity, provides critical insight into microalgal virus lineages, and shows that members of *Marnaviridae* extend beyond well-studied marine systems into freshwater ecosystems. Understanding the diversity and activity of these viruses is essential for clarifying how viral processes influence microbial community structure, phytoplankton dynamics, and nutrient fluxes across aquatic ecosystems.

## Materials and Methods

### Sample collection and sequencing

Metatranscriptomic datasets were generated from aquatic samples collected at five freshwater sites in southern Finland: Vantaanjoki river (60.2163° N, 24.9810° E) and the lakes Bodominjärvi (60.2419° N, 24.6583° E), Sääksjärvi (60.5113° N, 24.6575° E), Bakträsk (60.1875° N, 24.3628° E), and Luukinjärvi (60.3249° N, 24.6616° E), during June-August 2023 (Table S1). One coastal marine sample was collected from Hiroshima Bay in the Seto Inland Sea, Japan (34.2952° N, 132.4341° E), in March 2023. The Hiroshima Bay sample was included as an exploratory external comparison with the Finnish freshwater samples, not as a balanced marine-freshwater comparison.

Samples were concentrated from surface waters using a 20-µm mesh plankton net, thereby targeting the plankton-retained fraction and potentially underrepresenting free viral particles and viruses associated with pico- and nanoplankton. The resulting plankton pellet was transferred to a nearby laboratory within 30 min to 1 h of collection. The pellet was gently washed with sterile seawater to remove loose ambient debris while retaining plankton-associated microbiota and was subsequently stored at -80 °C prior to RNA extraction.

Total RNA was extracted using a QIAGEN RNA isolation kit (QIAGEN, Hilden, Germany) following a modified mechanical-disruption and lysis protocol optimized for unicellular algal matrices, as described by Charon *et al.*^12^. The riboPOOL rRNA Depletion Seawater kit (SiTools) was used for rRNA depletion of 1 µg (microgram) of total RNA. RNA concentration and integrity were assessed before library preparation using capillary electrophoresis on an Agilent 2100 Bioanalyzer (Agilent Technologies, Santa Clara, CA, USA). Metatranscriptomic library preparation with TruSeq Stranded Total RNA (Illumina) and paired-end sequencing were performed by the DNA Sequencing and Genomics Laboratory, Institute of Biotechnology, Helsinki Institute of Life Science (HiLIFE), University of Helsinki, Finland, using the AVITI platform with the AVITI High Output Cloudbreak Freestyle 2 x 150 kit (Element Biosciences). The library pool was loaded at 10 pM, and sequencing was performed in high-density mode.

### Read processing

Raw sequencing reads were initially assessed for base quality, adapter contamination, and sequence composition using FastQC v0.11.9^17^. Based on these quality control metrics, reads were trimmed using fastp v0.24.0^18^ with automatic paired-end adapter detection, removal of the first 10 bases from both reads, a minimum Phred quality score of 30 and a minimum post-trimming read length of 50 bp (‘--detect_adapter_for_pe -q 30 -l 50 --trim_front1 10 --trim_front2 10’). The second round of quality assessment with FastQC confirmed that only high-quality reads were retained for downstream analyses.

### Assembly

Ribosomal RNA (rRNA) reads were removed using SortMeRNA v4.3.7 with the default parameters^19^. Reads aligning to rRNA sequences with an E-value <0.001 were discarded to enrich messenger RNA (mRNA) transcripts prior to assembly. The remaining non-rRNA reads were assembled *de novo* into contigs with rnaSpades v4.0.0^20^ and assembly quality was evaluated using rnaQUAST v2.3.0^21^.

### Predicting and annotating viral operational taxonomic units (vOTUs)

Viral contigs were identified and processed using the Modular Viromics Pipeline (MVP) v1.1.5^22^. Assembled viral contigs ≥1 kb were first detected using geNomad v1.7.4 with database v1.7^23^. Sequence completeness and quality were evaluated with CheckV v1.0.3 using database v1.5^24^. Contigs were filtered to retain those that encoded at least one viral gene and displayed a viral-to-host gene ratio of at least 1:1.

Validated viral contigs were subsequently clustered into species-level vOTUs at a threshold of 95% average nucleotide identity (ANI) and 85% alignment coverage, as implemented in MVP. To quantify viral abundance across samples, non-rRNA reads were mapped to viral contigs using Bowtie2 v2.5.4^25^. Mapping results were summarized using SAMtools v1.21^26^ and CoverM v0.7.0^27^ into reads per kilobase of transcript per million mapped reads (RPKM) within MVP, and a horizontal coverage threshold of 50% was used to retain RPKM values for further analyses. Among predicted vOTUs, transposable elements (TEs) were identified using TEsorter v1.4.7^28^ with an E-value threshold of 0.001. Functional annotation of vOTUs was performed within MVP module 6. Additional gene function predictions for DNA vOTUs that were classified as prokaryotic viruses were obtained using Phold v0.2.0^29^.

### Viral host prediction

Putative bacterial and archaeal hosts for prokaryotic vOTUs were predicted using iPHoP v1.4.1^30^ with the Jun_2025_pub_rw database. Confidence score thresholds of 75 and 90 were applied for family- and genus-level assignments, respectively. When multiple host predictions were obtained, only the highest-confidence hit was retained.

For eukaryotic RNA vOTUs, putative hosts were predicted using RNAVirHost v1.0.5^31^. Viral order assignments from MVP were used where available; otherwise, the internal prediction tool of RNAVirHost was applied. Predictions labeled “high confidence”, or “assign” were retained. All predictions were cross-validated against ViralZone^32^, which resulted in the manual removal of two mispredictions, which represented a mismatch between MVP taxonomic assignment (*Leviviricetes*) and RNAVirHost predicted host (Chordata, Artiodactyla).

### Genomic comparisons for *Marnaviridae* vOTUs

ORFs within the two vOTUs that were classified as *Marnaviridae* by geNomad were predicted using the NCBI Open Reading Frame Finder (https://www.ncbi.nlm.nih.gov/orffinder/), with the minimum ORF length set to 600 nt, the standard genetic code, and nested ORFs excluded. Predicted ORF products were annotated using HHPred searches against the PDB_mmCIF70_20_Feb and SCOPe70_2.08 databases with a probability threshold of 95% and NCBI Conserved Domain searches against the CDD database with an E-value threshold of 0.001 (https://www.ncbi.nlm.nih.gov/Structure/cdd/wrpsb.cgi).

For phylogenetic analysis, ORFs of the two putative *Marnaviridae* vOTUs were translated into amino acid sequences using the ExPASy Translate tool^33^. Genome identifiers for classified members of the family *Marnaviridae* were obtained from the ICTV database (https://ictv.global/taxonomy), and corresponding RdRp-containing amino acid sequences were retrieved from NCBI^34^. For the phylogenetic tree including unclassified members of *Marnaviridae*, the genome identifiers were also collected from the ICTV database, and if there were multiple strains from the same virus species, only one strain was selected for the analysis. RdRp regions were identified with InterProScan^35^ by detecting domains annotated as RNA/DNA polymerases, and the corresponding regions were extracted with a custom Python script. RdRp sequences were aligned using the MAFFT v7.490 L-INS-i algorithm^36^ and trimmed with trimAl v1.2rev59^37^ using the automated1 mode. Maximum-likelihood phylogenetic inference was performed with IQ-TREE v2.0.7^38^, with ModelFinder^39^ used for substitution model selection and ultrafast bootstrap analysis^40^ used to assess branch support. The tree was rooted using RdRps of the *Dicistroviridae* family as an outgroup. Phylogenetic trees were visualized with iTOL v7.5.1^41^.

Pairwise genome comparisons between the *Marnaviridae* vOTUs were performed using Easyfig v2.2.5^42^ with BLASTn alignments generated at an E-value threshold of 1 × 10⁻⁵. To identify related reference sequences, *Marnaviridae* vOTUs were used as queries in BLASTn searches against the NCBI core_nt database and the IMG/VR v4 database^23^ using the same E-value threshold. NCBI core_nt hits with at least 80% query coverage were retained for downstream comparison. For IMG/VR hits, only high-quality uncultivated viral genomes (UViGs) were retained, and if a few retained UViGs represented the same vOTU, the longest UViG from each vOTU cluster was selected for visualization. Whole-genome comparisons between the recovered *Marnaviridae* vOTUs and retained NCBI or IMG/VR sequences were visualized with the Circoletto web server^43^ via BLASTn alignments at an E-value threshold of 1 × 10⁻⁵; with alignment ribbons color-coded according to absolute nucleotide identity percentages.

## Results

### Recovered vOTUs represented diverse RNA and DNA viruses

Initial MVP output included 449 vOTUs, of which 50 were recognized as TEs (Table S2), predominantly long terminal repeat retrotransposons (LTR-RTs). Thus, 399 vOTUs were retained for downstream analyses (Table S3). The recovered virome was largely composed of short (1-5 kb, 333 vOTUs, 84% of all vOTUs) fragmented genomes (Figure 1A), consistent with the complexity of environmental metatranscriptomic assemblies. CheckV analysis estimated 218 vOTUs (54.6%) as low-quality genomes, whereas 88 vOTUs (22%) were assigned as medium quality, while for 32 vOTUs (8%), the quality could not be confidently assessed (Figure 1B). Notably, 54 vOTUs showed 95-100% completeness, representing mostly RNA viruses and two DNA viruses. The longest genomes that were recovered (15-43 kbp) belonged to DNA viruses (see below for more details about DNA viruses), while the longest RNA vOTU was 13 kb in length.

**Figure 1:**
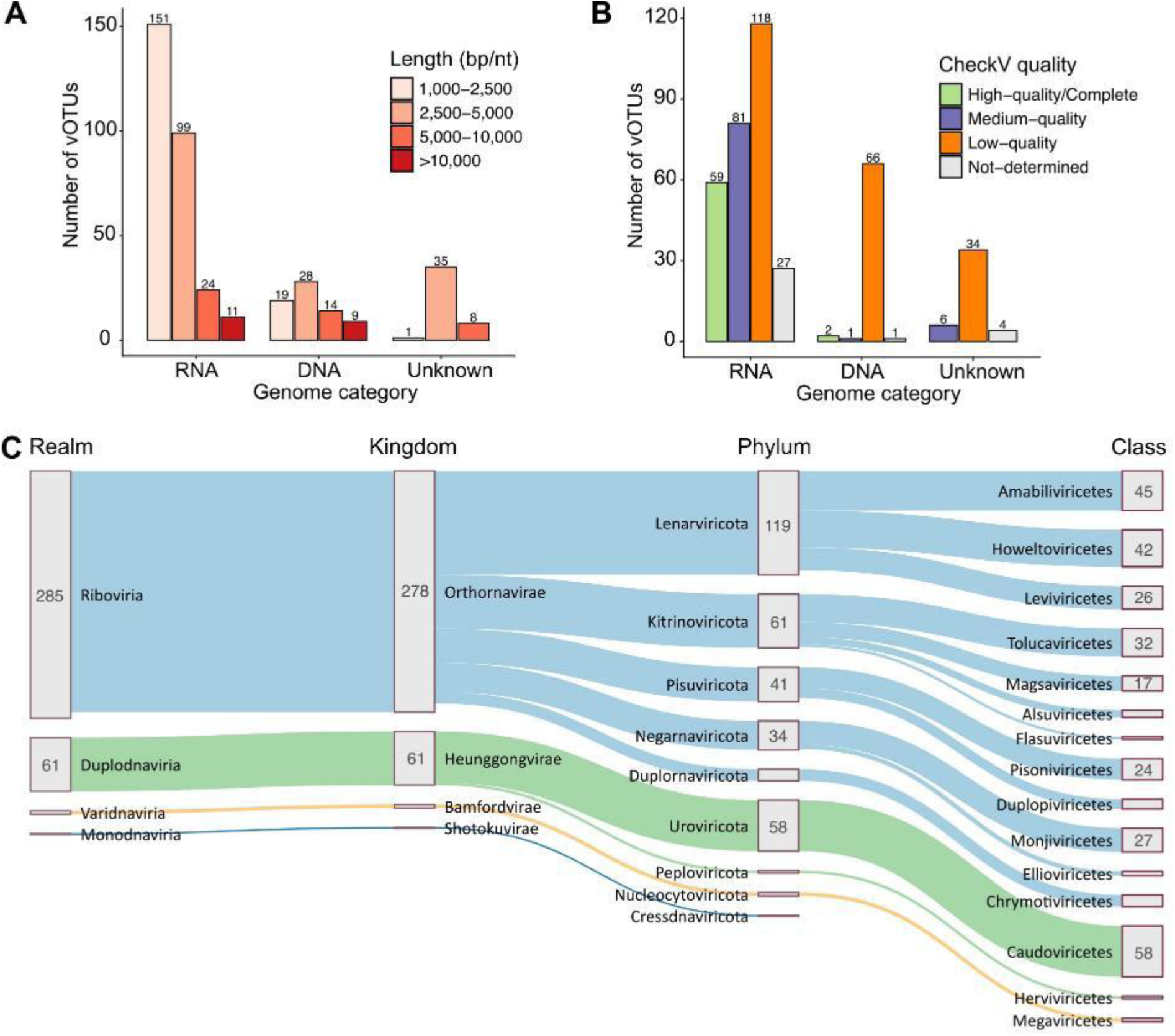
Overview of the 399 recovered viral operational taxonomic units (vOTUs). Distribution of the recovered vOTUs (A) by length and (B) across CheckV quality categories, vOTUs are grouped by genome type. (C) Taxonomic composition of classified vOTUs as predicted with geNomad. Unclassified vOTUs are not shown. Numbers indicate the total number of vOTUs assigned to each taxonomic group.

Taxonomic classification of vOTUs by geNomad revealed a virome overwhelmingly dominated by RNA viruses, particularly members of the realm *Riboviria* (285 vOTUs), whereas DNA viruses accounted for a substantially smaller fraction of classified sequences (70 vOTUs; Figure 1C). About 11% of vOTUs (44 vOTUs) could not be classified and remained of unknown genome types.

Within the RNA viruses, most vOTUs were classified within the phylum *Lenarviricota* (119 vOTUs), comprising the classes *Amabiliviricetes*, *Howeltoviricetes,* and *Leviviricetes* (Figure 1C). The second most represented RNA-virus phylum was *Kitrinoviricota* (61 vOTUs), approximately half of which were classified within the class *Tolucaviricetes* (32 vOTUs). Other predicted RNA-virus phyla included *Pisuviricota* (41 vOTUs), *Negarnaviricota* (34 vOTUs), and *Duplornaviricota* (13 vOTUs). Near-complete RNA vOTUs included representatives of the classes *Howeltoviricetes* (*Mitoviridae*), *Amabiliviricetes* (*Narnaviridae*), *Magsaviricetes* (*Nodaviridae*), *Tolucaviricetes* (*Tombusviridae*), *Alsuviricetes* (*Betaflexiviridae*), *Pisoniviricetes* (*Calicivirida*e, *Marnaviridae* and unclassified families), *Flasuviricetes* (*Flaviviridae*), *Monjiviricetes* (unclassified at the family level), and *Leviviricetes* (unclassified at the family level). These assignments indicate that the recovered sequences span diverse RNA-virus groups, although taxonomy alone does not confirm their hosts.

DNA viruses accounted for ∼17.5% (70 vOTUs) of all recovered vOTUs and were represented predominantly by prokaryotic dsDNA tailed viruses from the class *Caudoviricetes* (58 vOTUs). Among these, one vOTU classified within the family *Autographiviridae* was estimated as complete and was also the longest vOTU (42,579 bp) observed in our dataset. The other groups of DNA vOTUs included eukaryotic viruses from the phylum *Nucleocytoviricota* (5 vOTUs, four of which were classified as *Megaviricetes*) and *Peploviricota* (3 vOTUs within the family *Herpesviridae*). In addition, four ssDNA vOTUs were recovered, including three eukaryotic viruses from the phylum *Cressdnaviricota* and one complete *Microviridae* genome (phylum *Phixviricota*) (Figure 1).

To further assess potential host associations, host prediction was performed for eukaryotic RNA with the RNAVirHost tool and for all prokaryotic vOTUs with iPHoP (Table S3). RNAVirHost assigned putative hosts to 243 eukaryotic RNA vOTUs, with predictions dominated by Fungi, Invertebrate and Viridiplantae. When restricted to “assign” or high-confidence predictions, 152 eukaryotic RNA vOTUs retained host assignments, including 91 fungal, 43 invertebrate and 18 Viridiplantae-associated predictions. iPHoP assigned putative bacterial hosts to nine prokaryotic vOTUs: four vOTUs were linked to the phylum *Pseudomonadota* (the families *Enterobacteriaceae, Vibrionaceae, Pseudomonadaceae*), two to the phylum *Bacteroidota* (the families *Crocinitomicaceae* and *Bacteroidaceae*), and three vOTUs to the phylum *Patescibacteriota* (the families UBA4665, UBA1547, and UBA2103). Interestingly, the latter phylum includes ultra-small bacteria that have remained largely uncultivated and are widespread in diverse environments, especially groundwater systems^44^.

### Sample-specific viral profiles

The viral and TE profiles varied markedly among samples, with each environment showing a distinct dominant signal (Figure 2). TEs were detected across all Finnish samples but were entirely absent in the Japanese dataset. Lake Bakträsk was dominated by vOTUs classified as *Flaviviridae*, a family that includes positive-sense RNA viruses best known from animal hosts, which accounted for approximately three-quarters of the viral/TE %RPKM signal but was only weakly represented in the other datasets (Figure 2). By contrast, the Hiroshima Bay marine sample was dominated by *Mitoviridae***-**classified vOTUs, particularly unclassified *Caudoviricetes* and *Leviviricetes.* The Luukinjärvi and Bodominjärvi libraries contain a notable fraction of unclassified viral sequences. Bodominjärvi and Sääksjärvi showed high relative abundance of *Tombusviridae*, a family mainly comprising plant-associated positive-sense RNA viruses, suggesting a prominent aquatic plant- or algae-associated viral component in these samples.

**Figure 2:**
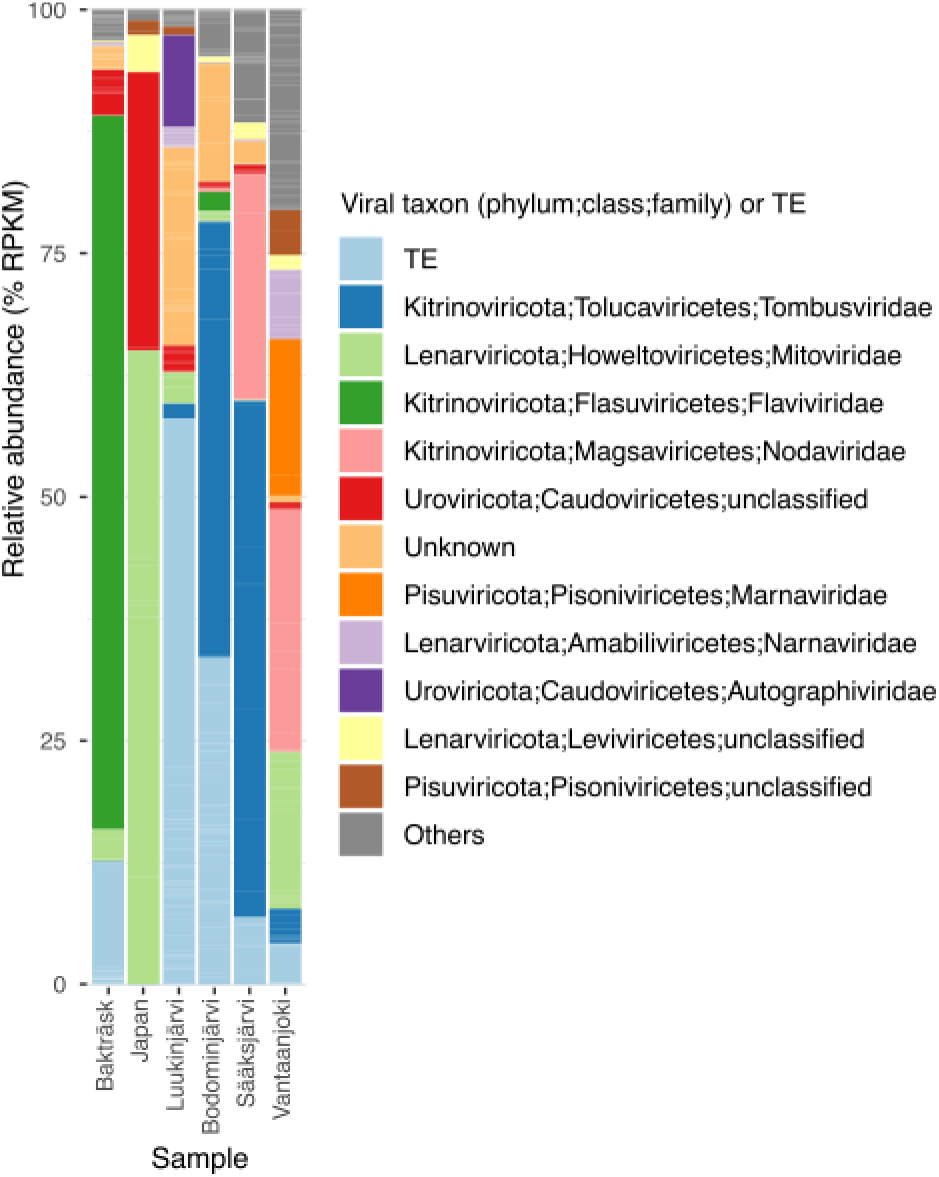
Relative abundance of viral taxa and TEs across samples. Stacked bar plot showing the relative abundance of viral taxa and TEs across samples, expressed as %RPKM (horizontal coverage of 50% applied). Each bar represents one sample, and colored segments indicate the proportional contribution of classified viral families, higher-level unclassified viral groups, and TEs. Colors correspond to the taxonomic groups shown in the legend on the right.

The Vantaanjoki river sample showed the most heterogeneous viral profile, yielding a high relative abundance of *Marnaviridae*, together with notable shares from vOTUs classified within *Nodaviridae*, *Mitoviridae*, *Narnaviridae,* and unclassified *Picornavirales/Pisoniviricetes* vOTUs (Figure 2). These groups suggest contributions from microeukaryote-associated, fungi-associated, and metazoa-associated RNA viruses within the same riverine environment. vOTUs classified as *Nodaviridae*, the family that includes viruses of insects and fish, were also prominent in Sääksjärvi, indicating that metazoa-associated viral signatures varied among the sampled environments.

### Transcriptionally active (DNA) vOTUs represented prokaryotic and eukaryotic viruses

DNA-virus sequences recovered from a metatranscriptome may indicate transcription of viral genes, but their detection alone does not demonstrate a productive infection. To characterize their predicted coding potential, eukaryotic DNA vOTUs were annotated with MVP/geNomad (Table S4), whereas prokaryotic DNA vOTUs were analysed using Phold (Table S5). The eukaryotic DNA-virus fraction included 11 vOTUs and was only partly resolved. Across these vOTUs, 16 ORFs were predicted, nine of which received functional annotations from geNomad. These included annotations comprised a limited set of functions compatible with DNA virus replication, genome maintenance or host interaction, including three putative viral replication proteins, two kinase-like proteins, a Rad2-like DNA excision-repair protein, a plasmid-replication-associated AAA-domain protein, an mRNA-capping enzyme/guanylyltransferase alpha subunit, and a UL97-like protein. These annotations indicate that some recovered sequences encode proteins compatible with DNA-virus functions, although the incomplete assemblies and limited annotations constrain functional interpretation.

The prokaryotic DNA-virus fraction was larger (59 vOTUs) and functionally better resolved than its eukaryotic counterpart. Across the prokaryotic DNA vOTUs, Phold assigned 394 functional annotations (Figure S1), of which 184 were classified as hypothetical or unknown proteins (46.7%). Among the resolved annotations, the largest category was DNA, RNA, and nucleotide metabolism (74 annotations; 18.8%), followed by virion head and packaging modules (44; 11.2%) and tail-associated structural functions (38; 9.6%). Connector proteins, lysis functions, auxiliary metabolic genes (AMGs) or host-takeover genes, integration/excision elements, and transcriptional regulators were less frequent. These annotations are consistent with bacteriophage-like genomic modules, while the large fraction of unknown proteins and fragmented assemblies limits functional interpretation.

The largest complete phage-like vOTU in the dataset (Figure S1) encoded genome-processing functions, including DNA polymerase, DNA primase/helicase, RNA polymerase, single-stranded DNA-binding or annealing proteins, a RusA-like Holliday-junction resolvase, endonuclease VII, exonuclease, DNA methyltransferase, integrase, and a transcriptional regulator. These genes occurred alongside predicted structural and packaging functions, including head-assembly and major-head protein, internal virion proteins, small and large terminase subunits, a head-tail adaptor, tail proteins, and tail-fiber proteins, as well as genes encoding lysis-associated amidase and endolysin-like proteins. Together, these annotations support its interpretation as a well-resolved bacteriophage-like genome. No host was predicted for A005_NODE_1; no inference about its actual host or infection dynamics can be made from the annotations alone.

The remaining *Caudoviricetes* vOTUs represented more compact or fragmented and functionally distinct phage modules (Table S5). For example, A005_NODE_87 was dominated by structural and genome-delivery annotations, including DNA ejection protein, head morphogenesis protein, head closure protein, major head protein, head scaffolding protein, portal protein, large terminase subunit, tail fiber protein, and major tail protein. Interestingly, it also encoded a ferredoxin, which was placed in the annotation category “moron, AMG and host takeover”. Conversely, A005_NODE_277 carried recombination, genome-maintenance and host-interaction annotations, including an Erf-like single-stranded DNA annealing protein, NinB/Orf recombination mediator, anti-restriction protein, integrase, endolysin, and DNA repair exonuclease. Together, these vOTUs indicate that the prokaryotic DNA virus fraction included phage genomes or genome fragments with structural, replication, recombination and host-associated modules, reflecting fragmentation of the recovered vOTUs, but also a mosaic nature of phage genomes^45^.

Abundance mapping across libraries provided additional ecological context for the DNA vOTUs. At a 50% horizontal coverage threshold, most prokaryotic DNA vOTUs were detected in one sample, but 15 vOTUs spanned a few samples (Figure S1). The longest complete *Caudoviricetes* vOTU mentioned above (A005_NODE_1) was detected in two samples out of six, and the other complete prokaryotic DNA vOTU, A001_NODE_172, was detected in all six samples. This latter contig was the only *Microviridae* representative in the prokaryotic DNA virus set. Phold annotations for this vOTU included terminase, which was manually removed as a likely misannotation, DNA replication initiation protein, major head protein, major spike protein, pilot protein for DNA ejection (initially labeled as “tail” category, which we manually re-labelled as “other”), and DNA replication protein in addition to a few hypothetical proteins with unknown functions (Table S5, Figure S1). A host prediction linked A001_NODE_172 to the genus *Escherichia,* although this should be treated as a putative host assignment rather than evidence of infection.

### Genomic architecture of two novel *Marnavirida*e vOTUs

After characterizing the DNA-virus sequences recovered from the metatranscriptomes, we next focused on two RNA vOTUs, A004_NODE_9 and A004_NODE_28, putatively classified within the family *Marnaviridae* by geNomad. Additional NCBI core_nt searches using unclassified *Picornavirales* vOTUs did not retrieve further Marnaviridae-like matches. The abundance analysis detected both vOTUs only in the Vantaanjoki river sample (log10(RPKM) values of 4.5 for A004_NODE_9 and 3.7 for A004_NODE_28.

A more detailed genome organization analysis of A004_NODE_9 and A004_NODE_28 (Figure 3) supported their taxonomic assignment within the family *Marnaviridae*. A004_NODE_9 was assessed as a complete viral genome by CheckV. This vOTU was 9.7 knt in length and contained two ORFs, which encoded replication- and capsid-associated proteins, as suggested by both HHPred and NCBI CDD searches. One of the ORFs encoded non-structural replication domains, including RNA helicase and RdRp, whereas the other one encoded classic picornavirus-like capsid domains (Figure 3). This organization is consistent with a dicistronic marnavirus-like genome, where replication and structural modules are encoded separately.

**Figure 3:**
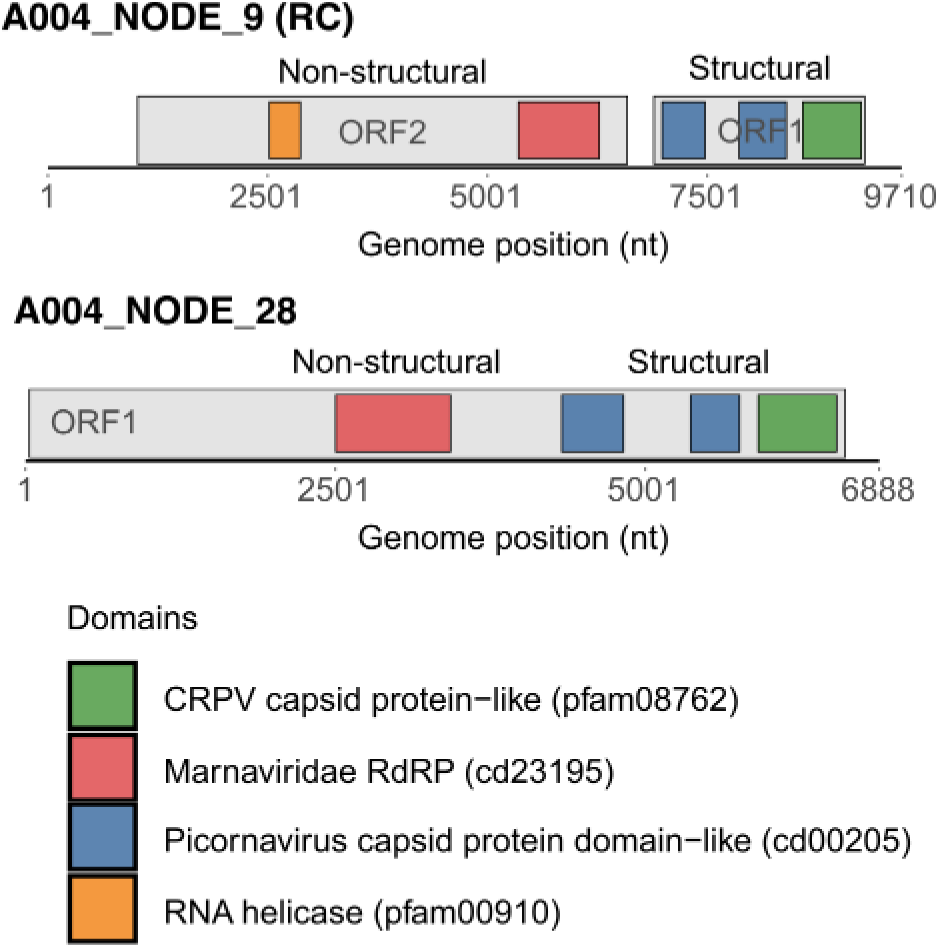
Predicted genome organization of the two *Marnaviridae* vOTUs identified in this study, A004_NODE_9 (shown as RC, reverse complement) and A004_NODE_28. ORFs were predicted using the standard genetic code. For A004_NODE_9, two ORFs were identified in the reverse-complemented sequence: ORF1 at nt 6892-9288 and ORF2 at nt 1022–6589. For A004_NODE_28, one long ORF was predicted from nt 32-6613. Conserved domains detected by NCBI CDD searches are shown as colored regions, with accession numbers indicated in brackets. HHPred searches independently supported the presence of picorna-like capsid protein and RNA-directed RNA polymerase (RdRp) signatures. CRPV, cricket paralysis virus.

A004_NODE_28 was 6.9 knt in length and assessed as 79%-complete. In contrast to A004_NODE_9, this vOTU encoded a single long ORF spanning most of the contig. HHPred and NCBI CDD searches identified RdRp and picornavirus-like capsid domains in distinct regions of the predicted polyprotein (Figure 3). The upstream placement of the RdRp relative to the capsid-associated domains preserves the replication-to-structural polarity expected for mono-partite *Marnaviridae* genomes.

### Phylogenetic Placement and the Candidate Genus “Vandarnavirus”

To investigate the taxonomic position of A004_NODE_9 and A004_NODE_28, we inferred a maximum-likelihood phylogeny from the conserved RNA-dependent RNA polymerase (RdRp) regions. Predicted ORFs were translated into amino acid sequences, and the RdRp regions were identified using InterProScan. The analysis was designed to assess their placement relative to classified *Marnaviridae* and related environmental sequences; phylogenetic placement alone does not establish host identity or formal taxonomic status.

In the RdRp phylogeny of classified *Marnaviridae,* A004_NODE_9 and A004_NODE_28 formed a strongly supported monophyletic lineage (ultrafast bootstrap support = 100) within the family-level grouping, but outside the currently recognized genera *Bacillarnavirus, Kusarnavirus, Labyrnavirus, Locarnavirus, Marnavirus, Salisharnavirus,* and *Sogarnavirus* (Figure 4). The new lineage recognized here, which we refer to as the Vantaanjoki lineage from hereafter, branched adjacent to, but remained distinct from, genus *Bacillarnavirus*, indicating that these two vOTUs are not assignable to any established *Marnaviridae* genus based on the RdRp phylogeny^16^. These results support recognition of a distinct environmental lineage, but formal genus-level classification requires evaluation under the relevant taxonomic criteria.

**Figure 4:**
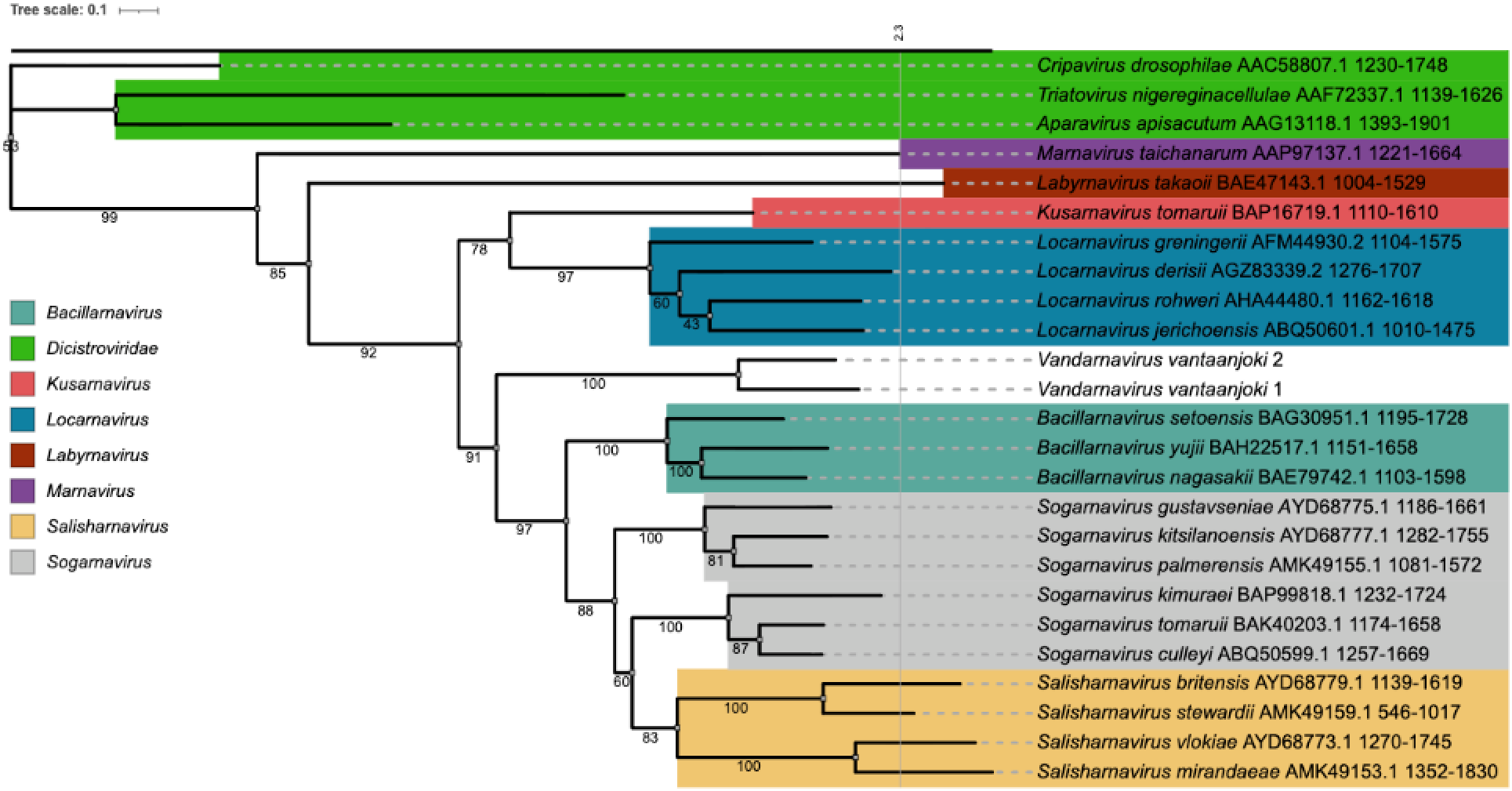
RdRp phylogeny of the two vOTUs proposed as the candidate genus “*Vandarnavirus*” vOTUs within *Marnaviridae*. Maximum-likelihood phylogeny inferred from the conserved RdRp region of A004_NODE_9 and A004_NODE_28, provisionally referred to as “*Vandarnavirus vantaanjoki1*” and “*Vandarnavirus vantaanjoki2*”, respectively, together with representative species of *Marnaviridae* recognized by the ICTV. Branch labels indicate ultrafast bootstrap support values. Recognized *Marnaviridae* genera are indicated by coloured ranges. The tree is rooted using representative *Dicistroviridae RdRp* as an outgroup. The scale bar represents amino-acid substitutions per site. GenBank identifiers and RdRp-region coordinates are shown where available. The tree was inferred under the LG+I+G4 model^46^.

An expanded RdRp analysis that included classified *Marnaviridae* representatives and additional *Marnaviridae*-like sequences listed by ICTV, yielded the same overall placement, with A004_NODE_9 and A004_NODE_28 remaining as a distinct Vantaanjoki lineage outside currently recognized genera. A004_NODE_9 and A004_NODE_28 again clustered together with maximal support and remained outside the currently recognized genera, within a broader environmental radiation that included *Sogarnavirus, Bacillarnavirus, Salisharnavirus,* and other picorna-like sequences. This broader context argues against an artefact of sparse taxon sampling and supports the interpretation that the Vantaanjoki sequences represent a distinct environmental lineage within *Marnaviridae* (Figure S2).

The genome organization and RdRp phylogeny support the provisional placement of A004_NODE_9 and A004_NODE_28 within *Marnaviridae* and indicate that they do not fall within any currently recognized genus in the analyses performed here. We therefore propose “*Vandarnavirus*” as a candidate genus name for this lineage, with the provisional candidate species names “*Vandarnavirus vantaanjoki1*” for A004_NODE_9 and “*Vandarnavirus vantaanjoki2*” for A004_NODE_28. These are proposed names for discussion and are not formal ICTV taxa.

### Viral sequences related to candidate “Vandarnavirus” vOTUs found from other environments

We compared the whole-genome sequences of A004_NODE_9 and A004_NODE_28 against the NCBI core_nt and IMG/VR v4 databases to identify related environmental sequences. A004_NODE_9 had two high-identity NCBI matches: *Picornavirales* sp. TC-PLV-28 MR233-17E/261^47^ genomic sequence (GenBank accession no: PP872584.1; 100% query coverage, 96.8% nucleotide identity) and Clirnapec virus 230 RdRp and capsid protein genes ^48^ (PQ161473.1; 100% coverage, 96.3% identity) (Figure 5A). The records were associated with metatranscriptomic studies: the TC-PLV-28 MR233-17E/261 sequence was recovered from river water collected in Berlin, Germany^47^, and Clirnapec virus 230 sequence was reported from an RNA virome associated with mussels grown in river water in Virginia, USA^48^. We therefore describe them as environmental picorna-like viral sequences rather than cultured isolates or metagenome-assembled genomes (MAGs).

**Figure 5:**
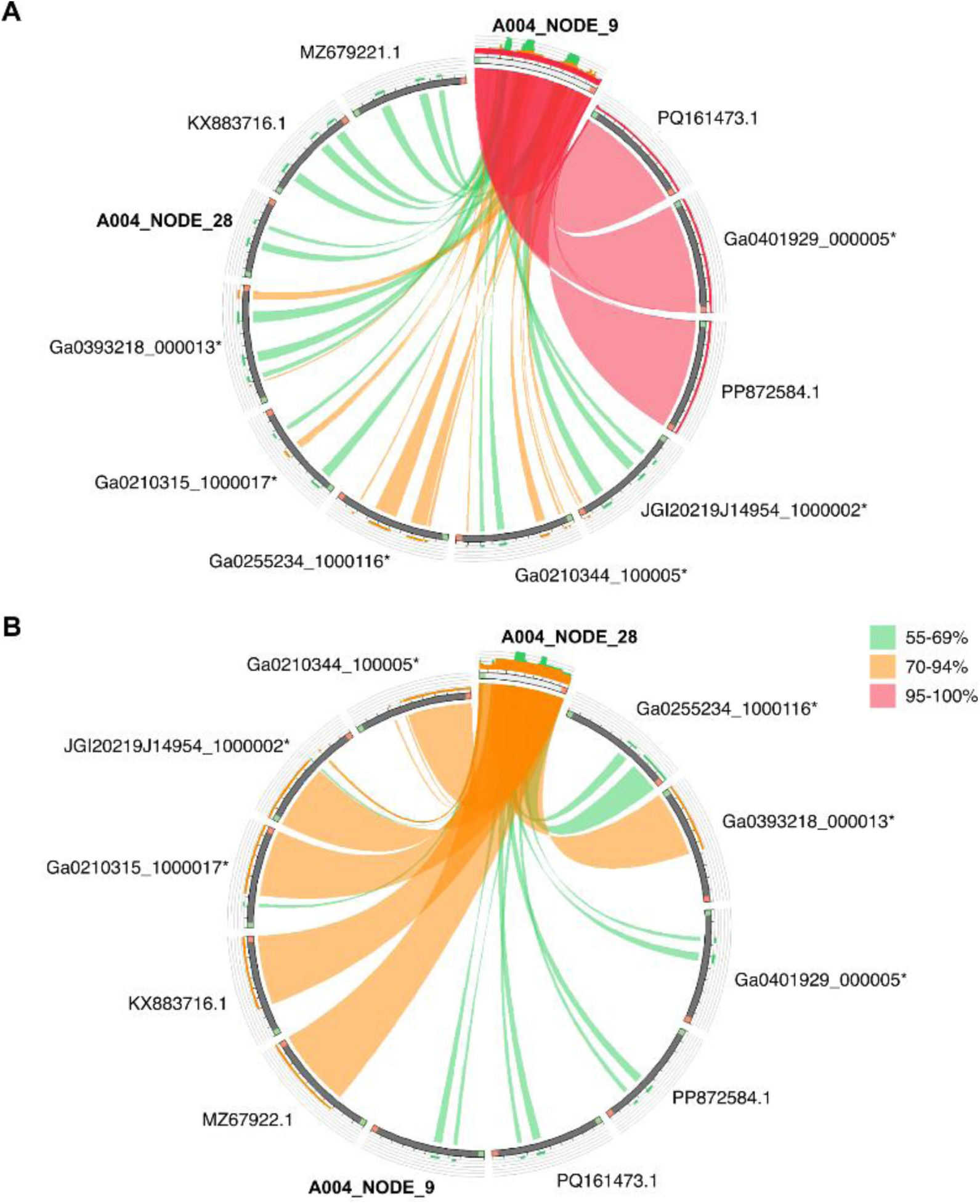
Whole-genome comparisons of the candidate genus “*Vandarnavirus*” vOTUs with related environmental viral sequences. Circoletto plots compare A004_NODE_9 (A) and A004_NODE_28 (B) (highlighted with bold) with selected NCBI core_nt records and high-quality uncultivated viral genomes (UViGs) from IMG/VR v4 (UViGs are marked with asterisks). The NCBI core_nt records represent environmental sequences reported in metatranscriptomic or RNA-virome studies, while the IMG/VR entries are UViGs. NCBI sequences are labelled by GenBank accession numbers and IMG/VR sequences by scaffold identifiers. For IMG/VR, the longest UViG representing each vOTU cluster was selected for visualization. BLASTn matches were visualized using an E-value threshold of 1 × 10^-^⁵ with ribbons coloured by nucleotide identity. The identity ranges are 63.4-100% for A004_NODE_9 and 65.5-77.0% for A004_NODE_28.

A004_NODE_28 was more divergent from the available NCBI core_nt references. Its closest NCBI core_nt matches were *Marnaviridae* sp. 373R-k141_428865^49^ hypothetical protein gene (GenBank accession no: MZ679221.1; 99% query coverage, 77.3% nucleotide identity) and Sanxia picorna-like virus 7 ^50^ hypothetical protein gene (KX883716.1; 88% coverage, 76.4% identity) (Figure 5B). The former sequence was recovered from terrestrial RNA viromes in China^49^, while the latter one was obtained from invertebrate RNA viromes^50^. Interestingly, Sanxia picorna-like virus 7 has previously been placed among divergent marnavirus-like sequences in the analyses of microalgal RNA virus diversity^12^. These records were derived from environmental RNA-virus datasets, including terrestrial and invertebrate-associated RNA viromes. Their database labels were retained only as identifiers; the sequences should not be described as cultured isolates unless the source records independently establish isolation in culture.

IMG/VR searches extended the comparison beyond named NCBI records (Table S6). For A004_NODE_9, 17 high-quality uncultivated viral genome (UViG) records representing three IMG/VR vOTU clusters were recovered. These included UViGs from environmental metatranscriptomes from surface sediment at Twitchell Island in the Sacramento–San Joaquin Delta, freshwater, soil, and wetland datasets from Ohio and Virginia, including Great Miami River, Stroubles Creek, Old Woman Creek estuary and Old Woman Creek Reserve datasets, and an estuarine sediment metatranscriptome from the Columbia River estuary, Washington. For A004_NODE_28, 21 high-quality IMG/VR UViG records representing five IMG/VR vOTU clusters were recovered (Table S6). These included the Twitchell Island surface-sediment cluster also detected for A004_NODE_9, salt-marsh sediment from Massachusetts, freshwater microbial metatranscriptomes from the Columbia River region, and Columbia River estuarine sediment and water. These matches indicate related environmental sequences across several habitat types, but they do not establish host range, activity, or ecological function.

Circoletto comparisons showed broad alignments between A004_NODE_9 and selected NCBI/IMG/VR representatives, with nucleotide identities ranging from 63.4% to 100% (Figure 5A). A004_NODE_28 showed lower-identity similarity to available sequences, with nucleotide identities ranging from 65.5% to 77.0% (Figure 5B), consistent with its more distant NCBI matches. Genome-wide comparison of database hits for the two *Marnaviridae* vOTUs reflected their divergence from each other, as the hits retrieved for each vOTU showed higher similarity to their respective query than to the other vOTU. These values describe similarity among the selected sequences and should not be interpreted as evidence of equivalent ecological roles or host associations.

## Discussion

Freshwater ecosystems represent a vast but uniquely dynamic and underexplored reservoir of viral diversity^51^. These systems support dynamic microbial communities that respond rapidly to environmental variability^52^, yet the viruses associated with them remain poorly described. This gap is especially relevant for freshwater phytoplankton and their viral pathogens. Coastal microalgal habitats likewise harbour productive microbial communities^53^, but the diversity and activity of their viruses remain insufficiently resolved. Recent work on seaweed and microalgal holobionts has similarly emphasized that viruses are an overlooked component of these host-associated microbial systems^14^.

To characterize viral sequences across contrasting aquatic habitats, we sampled freshwater systems in Finland and included a geographically distant marine sample from Hiroshima Bay, Japan. The Finnish freshwater sites represent the primary focus of this study, whereas the Hiroshima Bay sample was included as an exploratory external comparison. Because the sampling design was not balanced between freshwater and marine environments, this comparison should not be interpreted as a formal assessment of differences between the two environments.

The recovered RNA-virus sequences represented several taxonomic groups, including vOTUs assigned to *Mitoviridae* and *Narnaviridae*, which are often associated with fungi; *Tombusviridae*, which includes plant-associated viruses; *Nodaviridae,* which includes viruses of animals and protists; and *Leviviricetes*, which includes prokaryotic RNA viruses^54^. These family- or class-level associations provide context but do not identify the actual hosts of the individual vOTUs. The host predictions were computational, and no host transcriptomic or culture-based evidence was used here to confirm specific virus–host interactions.

Environmental RNA virome studies have greatly expanded known RNA virus diversity, including picorna-like and protist-associated lineages^55^, in line with large-scale global surveys that continue to highlight a massive, uncharacterized “dark matter” of aquatic RNA viruses^50,55,56^. Still, many recovered sequences lack close cultured representatives^13^. Moreover, RNA viromes are often challenging to produce complete viral genomes due to the segmented nature of some groups of RNA viruses^57^. In our dataset, many vOTUs were short, fragmentary, or only broadly classified, especially among eukaryote-associated RNA viruses. Therefore, biological interpretation remained limited by incomplete taxonomic resolution and sparse virus-host information. Still, 53.3% of RNA vOTUs received *in silico* host predictions to eukaryotic hosts. Future work, such as single-cell or population-resolved host transcriptomics, would be needed to resolve virus-host interactions in more detail.

Transposable-element-associated contigs contributed substantially to some sample profiles, particularly in Luukinjärvi and Bodominjärvi. Most were classified as long terminal repeat (LTR) retrotransposons and lacked meaningful NCBI core_nt matches, suggesting possible origins in poorly represented environmental eukaryotes. These elements may derive from microalgae or other freshwater eukaryotes, since LTR retrotransposons are abundant in e.g. diatom genomes and have been linked to genome plasticity and stress-associated variation in natural populations^58^. In aquatic environments, environmental stress or shifting nutrient levels may trigger the activation of mobile elements in wild phytoplankton^58^. We therefore interpret these contigs as expressed mobile-element sequences and do not assign them to a particular host lineage.

An additional limitation is the 20-µm mesh used for surface-water concentration. This approach targets the plankton-retained fraction and may underrepresent viruses associated with pico- and nanoplankton, as well as free viral particles that pass through the mesh. Consequently, the viral diversity characterized here should not be interpreted as representing the complete aquatic or microalgal virosphere. Rather, the results describe viral sequences recovered from the plankton-retained fraction of the sampled surface waters. Intracellular RNA viruses and actively transcribed DNA and RNA viruses associated with microeukaryotic, and bacterial cells retained by the sampling procedure may be enriched in the resulting metatranscriptomes. In our dataset, relative-abundance profiles differed among the six samples: Flaviviridae-like sequences were prominent in Bakträsk, Mitoviridae and prokaryotic-virus sequences contributed to the Hiroshima Bay profile, unclassified viral groups were prominent in Luukinjärvi, and Tombusviridae-like sequences were abundant in Bodominjärvi and Sääksjärvi. The Vantaanjoki profile contained a mixture of RNA-virus groups and the only detections of A004_NODE_9 and A004_NODE_28 in the updated analysis. However, these observations are descriptive and more replicates would be needed to assess whether the observed patterns represent ecologically distinct communities in the sampled environments. Future studies incorporating systematically collected physicochemical measurements alongside replicated sampling would allow viral community patterns to be examined in relation to environmental gradients. Specifically, this approach would enable the evaluation of associations between viral community composition and key environmental variables, such as temperature, pH, dissolved oxygen, chlorophyll-a, and nutrient concentrations.

While it remains difficult to assess the activity of the detected RNA viruses due to their various replication strategies, the DNA viral fraction recovered in the analysed metatranscriptomes suggested active viral infections. Eukaryotic DNA vOTUs were few and poorly annotated, whereas prokaryotic DNA vOTUs encoded recognizable phage modules involved in nucleotide metabolism, virion morphogenesis, genome packaging, integration, and lysis. The presence of a *Microviridae* vOTU in all six samples suggests that ssDNA phages may be recurrent across these libraries, consistent with the broad environmental distribution of microviru*s*-like phages^30^, but the predicted *Escherichia* host remains provisional and should not be taken as evidence of infection *in situ*. Likewise, other host links reported here should be considered as predictions only, including three *Caudoviricetes* vOTUs linked to the phylum *Patescibacteriota* (also known as Candidate Phyla Radiation, CPR). This phylum has been recognized mainly through (meta)genomic studies with only a few available co-cultures, suggesting an epibiotic lifestyle in these ultrasmall bacteria with reduced genomic repertoire^59^. Furthermore, single-cell genomics revealed that *Patescibacteriota* may be closely associated with microeukaryotes^60^. Extremely little is known about *Patescibacteriota*-associated viruses and how they interact with their hosts^61^. Despite their ultrasmall cell size and reduced metabolic capacity, *Patescibacteriota* can support at least limited phage infection, as shown by the recent experimental work^62^. Detecting transcriptionally active viruses putatively linked to *Patescibacteriota* in this study complements metagenomic studies that spotted uncultured phages linked to *Patescibacteriota* in groundwater and deep subsurface environments^63,64^ and further highlights the presence and activity of such viruses in freshwater environments.

This study focused further on two RNA vOTUs putatively assigned to *Marnaviridae*. The family belongs to *Picornavirales* and includes +ssRNA viruses associated with unicellular eukaryotes, including some algae^16^. In our data, A004_NODE_9 encoded a near-complete dicistronic genome with separate replication and capsid modules, whereas A004_NODE_28 was shorter but retained RdRp and picornavirus-like capsid domains. These features strongly support their assignment within *Marnaviridae*. Moreover, the RdRp phylogeny placed these two vOTUs in a distinct, strongly supported lineage within *Marnaviridae*, outside all currently recognized genera. Together with their genome organization, this suggests their provisional designation as members of a candidate for the new genus, “Vandarnavirus”. We propose to classify A004_NODE_9 and A004_NODE_28 as candidate species *Vandarnavirus vantaanjoki1 and Vandarnavirus vantaanjoki2*, respectively, within this new candidate genus. This proposal remains provisional pending formal ICTV evaluation.

The two candidates for marnavirus vOTUs differed in their relationship to known viral sequence space. While A004_NODE_9 was closely related to environmental picorna-like viral genomes, including a viral genome from the Teltow Canal, Berlin^47^, A004_NODE_28 shared only moderate similarity with environmental picorna-like sequences. IMG/VR comparisons further showed that relatives of both vOTUs occur in freshwater, wetland, saltmarsh, and estuarine metatranscriptomes. The broad distribution shows that while “Vandarnavirus” stands out as a unique lineage, it belongs to a widespread aquatic viral cluster. This fits a familiar pattern in global marine and freshwater surveys, where certain *Marnaviridae* clades routinely cross salinity gradients and geographic boundaries^65^. These relationships place the Vantaanjoki sequences within a broader environmental pool of marnavirus-like RNA viruses rather than representing isolated, localized genotypes.

In summary, the metatranscriptomes contained diverse viral and mobile-element sequences with tentative taxonomic links to fungal-, plant-, algal-, metazoan-, and prokaryote-associated groups. The principal finding is the recovery of two Marnaviridae-like vOTUs from Vantaanjoki with complementary genome-organization features and a distinct RdRp phylogenetic placement. Their provisional grouping as “Vandarnavirus” provides a testable taxonomic hypothesis and genomic targets for future studies. Host identity, infection dynamics, and effects on phytoplankton or nutrient cycling were not directly measured and remain questions for future work.

## Conclusion

Metatranscriptomic analysis recovered diverse viral sequences across five Finnish freshwater sites and one exploratory marine sample from Hiroshima Bay, Japan. The dataset included 399 vOTUs, of which 285 were classified as RNA viruses, alongside DNA-virus sequences with predicted viral functions. Two Marnaviridae-like vOTUs detected only in the Vantaanjoki river sample formed a distinct RdRp lineage outside currently recognized genera. We provisionally propose the candidate genus name “*Vandarnavirus*” for this lineage, pending formal taxonomic evaluation. Because sampling used a 20-µm plankton net, the results represent the retained plankton-associated fraction and may underrepresent smaller plankton and free virions. Future work combining broader size fractions, environmental metadata, and host-resolved experiments will be needed to establish host associations and evaluate the ecological significance of these viruses.

## Supporting information

Algal vOTUs ms supplementary tables

## Acknowledgements

We sincerely thank the Kone Foundation (Koneen Säätiö, Grant No. 202202610 to RS and Grant No. 201904346 to TD) for the generous financial support of this research. HMO was supported by the University of Helsinki and the Research Council of Finland by funding for FINStruct and Instruct Centre FI, part of Biocenter Finland and Instruct-ERIC. We also gratefully acknowledge the DNA Sequencing and Genomics Laboratory (BIDGEN) for their technical support and high-throughput sequencing services. We thank Keiko Hamada for helping in sample collection from Japan.

Anthropic Claude Sonnet 4.6 was used for developing R codes; these outputs were verified by TD.

## Data availability

Raw reads are available from the European Nucleotide Archive (ENA) under the project PRJEB114339. TEs and vOTUs sequences are available from Figshare under DOI:10.6084/m9.figshare.32620143 and DOI:10.6084/m9.figshare.32620080, respectively.

## Conflicts of Interest

The authors declare no conflict of interest.

## Supplementary Materials

**Figure S1:**
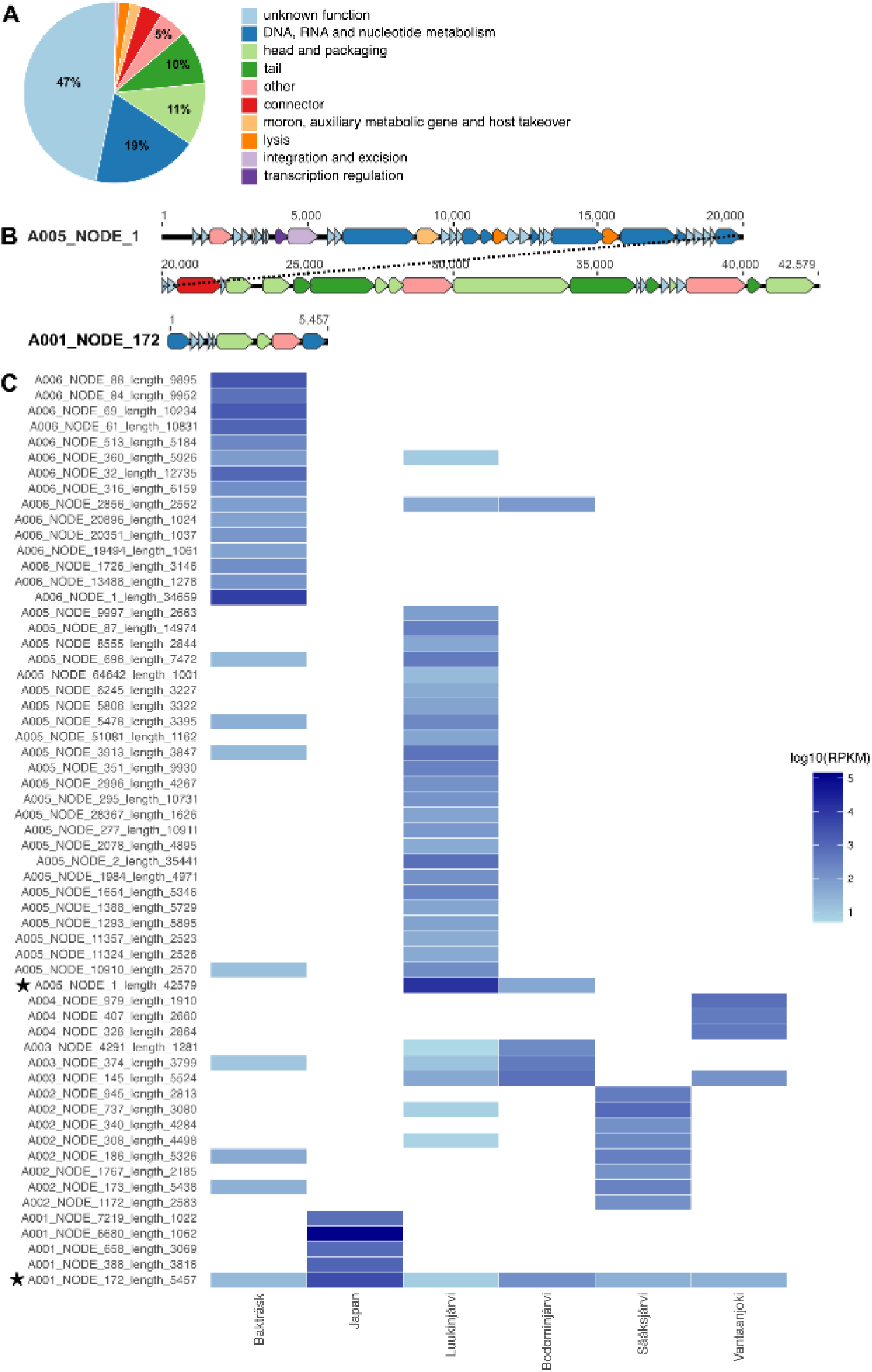
Functional annotation and sample distribution of transcriptionally detected prokaryotic DNA vOTUs. (A) Phold functional predictions for all prokaryotic DNA vOTUs. (B) Genome visualizations for complete prokaryotic DNA vOTUs: A005_NODE_1, putatively classified within the family *Autographiviridae* (order *Caudoviricetes*), and A001_NODE_172, putatively classified as *Microviridae* (*Malgrandaviricetes*). ORFs are shown as arrows. Note that the A005_NODE_1 genome is shown in two lines. Color-coding for ORF product functions is the same as in (A). (C) Distribution of prokaryotic DNA vOTUs across samples, based on mapping at a 50% horizontal coverage threshold. vOTUs that are shown in (B) are marked with stars in (C).

**Figure S2:**
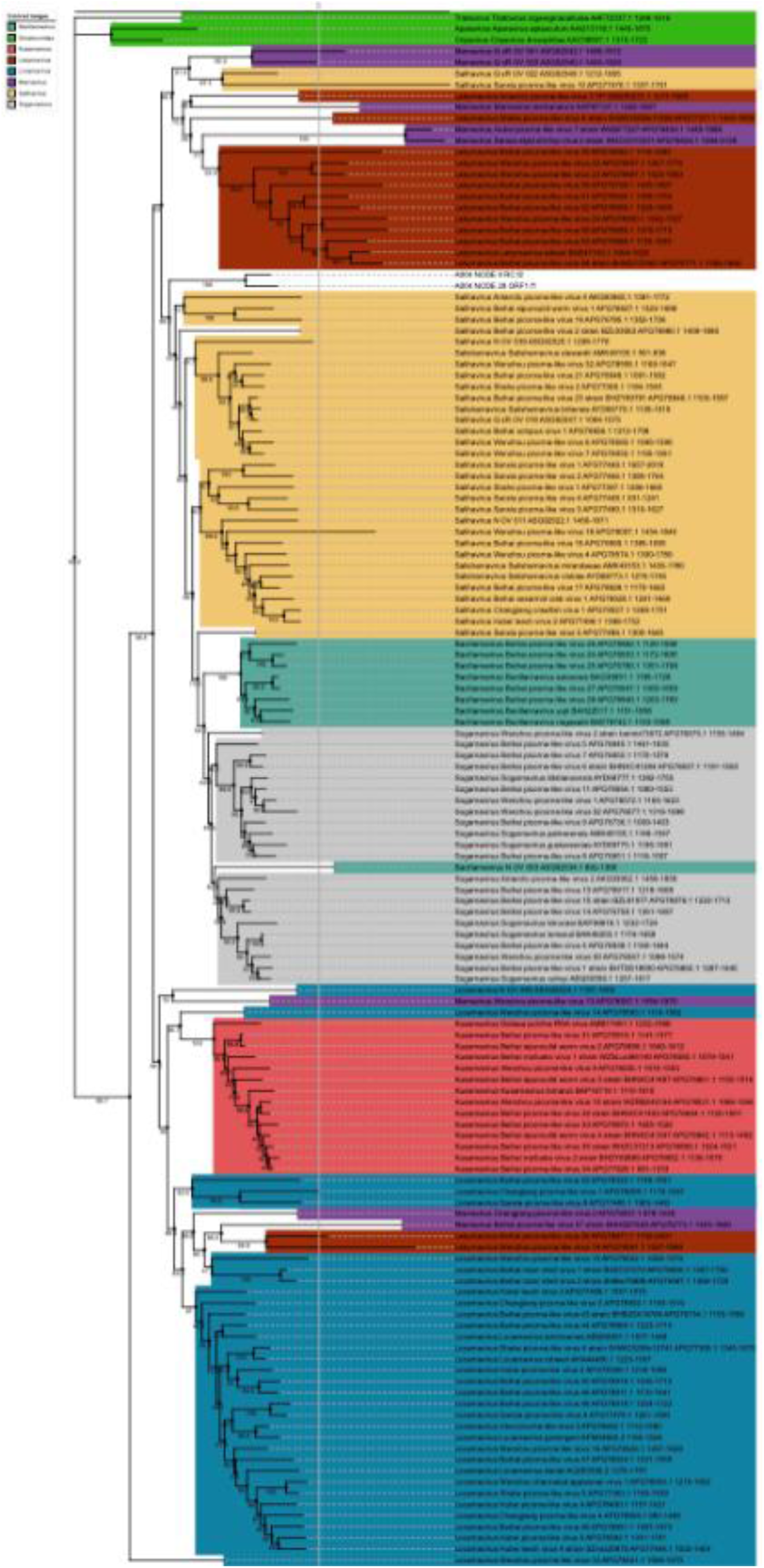
Expanded RdRp phylogeny of candidate “Vandarnavirus” vOTUs and environmental marnavirus-like sequences.

Table S1: Read-processing and transcriptome-assembly statistics for the six aquatic metatranscriptomic libraries. Summary of sequencing depth, read processing, ribosomal RNA recovery, and *de novo* assembly statistics for all samples analyzed in this study. Reported metrics include the total number of raw sequencing reads generated, accession numbers for raw read datasets, reads retained following quality trimming, percentage of SSU rRNA read pairs relative to total trimmed read pairs, total number of assembled transcripts, transcript N50, average transcript length, and length of the longest assembled transcript.

Table S2: Classification of transposable elements identified using TEsorter. Summary of transposable elements (TEs) detected across assembled contigs using TEsorter. Reported annotations include contig identity, transposable element order, superfamily classification, phylogenetic clade assignment, structural completeness, strand orientation, and conserved protein domains detected within each element

Table S3: vOTUs retained after transposable element removal. Catalogue of the 399 viral operational taxonomic units retained for downstream analyses, including genome quality, completeness, taxonomic assignment, genome type, host type and predicted host information where available.

Table S4: Gene-level MVP predictions for transcriptionally detected eukaryotic DNA vOTUs recovered from freshwater metatranscriptomes.

Table S5: Gene-level Phold predictions for transcriptionally detected prokaryotic DNA vOTUs, including functional categories related to phage replication, structure, packaging, integration and lysis.

Table S6: IMG/VR relatives of *Vantaanjoki Marnaviridae* vOTUs. High-quality IMG/VR v4 UViG hits retrieved using A004_NODE_9 and A004_NODE_28 as BLASTn queries. IMG/VR vOTU clusters are indicated to distinguish individual UViGs from species-level groups.

## Notes

### Competing Interest Statement

The authors have declared no competing interest.

## References

1. Falkowski, P. G., Katz, M. E., Knoll, A. H., Quigg, A., Raven, J. A., Schofield, O., & Taylor, F. J. R. (2004). The Evolution of Modern Eukaryotic Phytoplankton. Science, 305(5682), 354–360. 10.1126/science.1095964

2. Sadeghi, M., Tomaru, Y., & Ahola, T. (2021). RNA Viruses in Aquatic Unicellular Eukaryotes. Viruses, 13(3), 362. 10.3390/v13030362

3. Loos, L. M. van der, Eriksson, B. K., & Salles, J. F. (2019). The Macroalgal Holobiont in a Changing Sea. Trends in Microbiology, 27(7), 635–650. 10.1016/j.tim.2019.03.002

4. Zhao, J., Nair, S., Zhang, Z., Wang, Z., Jiao, N., & Zhang, Y. (2024). Macroalgal virosphere assists with host–microbiome equilibrium regulation and affects prokaryotes in surrounding marine environments. The ISME Journal, 18(1), wrae083. 10.1093/ismejo/wrae083

5. Coy, S. R., Gann, E. R., Pound, H. L., Short, S. M., & Wilhelm, S. W. (2018). Viruses of Eukaryotic Algae: Diversity, Methods for Detection, and Future Directions. Viruses, 10(9), 487. 10.3390/v10090487

6. Oleynik, G. N., Yurishinets, V. I., & Starosila, Ye. V. (2012). Viruses in Aquatic Ecosystems: Distribution and Ecological Role (a Review). Hydrobiological Journal, 48(5), 69–75. 10.1615/hydrobj.v48.i5.60

7. Robinson, D., Morgan-Kiss, R. M., Wang, Z., & Takacs-Vesbach, C. (2024). Antarctic lake viromes reveal potential virus associated influences on nutrient cycling in ice-covered lakes. Frontiers in Microbiology, 15, 1422941. 10.3389/fmicb.2024.1422941

8. Rosenwasser, S., Ziv, C., Creveld, S. G. van, & Vardi, A. (2016). Virocell Metabolism: Metabolic Innovations During Host–Virus Interactions in the Ocean. Trends in Microbiology, 24(10), 821–832. 10.1016/j.tim.2016.06.006

9. Edwards, K. F., McBeain, K. A., Schvarcz, C. R., & Steward, G. F. (2025). Large, double-stranded DNA viruses tend to suppress phytoplankton populations more effectively than small viruses of diverse genome type. Limnology and Oceanography Letters, 10(5), 745–753. 10.1002/lol2.70035

10. Chin, P.-J., Tsou, J. H., Armstrong, A., Deneyer, N., Zanda, V., Ayama-Canden, S., Colinet, A.-S., Fuentes, S. M., Korokhov, N., Lambert, C., Lavorgna, A., Noll, M., Olgiati, S., Protz, M., Shaid, S., Sohrabi, A., Whiteman, M., & Khan, A. S. (2025). Evaluation of high-throughput sequencing for replacing the conventional adventitious virus detection assays used for biologics. Npj Vaccines, 11(1), 28. 10.1038/s41541-025-01351-2

11. Pérez-Losada, M., Arenas, M., Galán, J. C., Bracho, M. A., Hillung, J., García-González, N., & González-Candelas, F. (2020). High-throughput sequencing (HTS) for the analysis of viral populations. Infection, Genetics and Evolution, 80, 104208. 10.1016/j.meegid.2020.104208

12. Charon, J., Kahlke, T., Larsson, M. E., Abbriano, R., Commault, A., Burke, J., Ralph, P., & Holmes, E. C. (2022). Diverse RNA Viruses Associated with Diatom, Eustigmatophyte, Dinoflagellate, and Rhodophyte Microalgae Cultures. Journal of Virology, 96(20), e00783–22. 10.1128/jvi.00783-22

13. Wolf, Y. I., Silas, S., Wang, Y., Wu, S., Bocek, M., Kazlauskas, D., Krupovic, M., Fire, A., Dolja, V. V., & Koonin, E. V. (2020). Doubling of the known set of RNA viruses by metagenomic analysis of an aquatic virome. Nature Microbiology, 5(10), 1262–1270. 10.1038/s41564-020-0755-4

14. Kolundžija, S., Cheng, D.-Q., & Lauro, F. M. (2022). RNA Viruses in Aquatic Ecosystems through the Lens of Ecological Genomics and Transcriptomics. Viruses, 14(4), 702. 10.3390/v14040702

15. Abbasi, M., & Alam, M. (2025). Understanding the roles of viruses as key players in environmental dynamics and ecosystem functioning. Discover Viruses, 2(1), 10. 10.1007/s44370-025-00015-y

16. Lang, A. S., Vlok, M., Culley, A. I., Suttle, C. A., Takao, Y., Tomaru, Y., & Consortium, I. R. (2021). ICTV Virus Taxonomy Profile: Marnaviridae 2021. The Journal of General Virology, 102(8), 001633. 10.1099/jgv.0.001633

17. Brandine, G. de S., & Smith, A. D. (2020). Falco: high-speed FastQC emulation for quality control of sequencing data. F1000Research, 8, 1874. 10.12688/f1000research.21142.2

18. Chen, S., Zhou, Y., Chen, Y., & Gu, J. (2018). fastp: an ultra-fast all-in-one FASTQ preprocessor. Bioinformatics, 34(17), i884–i890. 10.1093/bioinformatics/bty560

19. Kopylova, E., Noé, L., & Touzet, H. (2012). SortMeRNA: fast and accurate filtering of ribosomal RNAs in metatranscriptomic data. Bioinformatics, 28(24), 3211–3217. 10.1093/bioinformatics/bts611

20. Bushmanova, E., Antipov, D., Lapidus, A., & Prjibelski, A. D. (2019). rnaSPAdes: a de novo transcriptome assembler and its application to RNA-Seq data. GigaScience, 8(9), giz100. 10.1093/gigascience/giz100

21. Bushmanova, E., Antipov, D., Lapidus, A., Suvorov, V., & Prjibelski, A. D. (2016). rnaQUAST: a quality assessment tool for de novo transcriptome assemblies. Bioinformatics, 32(14), 2210–2212. 10.1093/bioinformatics/btw218

22. Coclet, C., Camargo, A. P., & Roux, S. (2024). MVP: a modular viromics pipeline to identify, filter, cluster, annotate, and bin viruses from metagenomes. mSystems, 9(10), e00888–24. 10.1128/msystems.00888-24

23. Camargo, A. P., Roux, S., Schulz, F., Babinski, M., Xu, Y., Hu, B., Chain, P. S. G., Nayfach, S., & Kyrpides, N. C. (2024). Identification of mobile genetic elements with geNomad. Nature Biotechnology, 42(8), 1303–1312. 10.1038/s41587-023-01953-y

24. Nayfach, S., Camargo, A. P., Schulz, F., Eloe-Fadrosh, E., Roux, S., & Kyrpides, N. C. (2021). CheckV assesses the quality and completeness of metagenome-assembled viral genomes. Nature Biotechnology, 39(5), 578–585. 10.1038/s41587-020-00774-7

25. Langmead, B., & Salzberg, S. L. (2012). Fast gapped-read alignment with Bowtie 2. Nature Methods, 9(4), 357–359. 10.1038/nmeth.1923

26. Li, H., Handsaker, B., Wysoker, A., Fennell, T., Ruan, J., Homer, N., Marth, G., Abecasis, G., Durbin, R., & Subgroup, 1000 Genome Project Data Processing. (2009). The Sequence Alignment/Map format and SAMtools. Bioinformatics, 25(16), 2078–2079. 10.1093/bioinformatics/btp352

27. Aroney, S. T. N., Newell, R. J. P., Nissen, J. N., Camargo, A. P., Tyson, G. W., & Woodcroft, B. J. (2025). CoverM: read alignment statistics for metagenomics. Bioinformatics, 41(4), btaf147. 10.1093/bioinformatics/btaf147

28. Zhang, R.-G., Li, G.-Y., Wang, X. L., Dainat, J., Wang, Z. X., Ou, S., & Ma, Y. (2022). TEsorter: An accurate and fast method to classify LTR-retrotransposons in plant genomes. Horticulture Research, 9, uhac017. 10.1093/hr/uhac017

29. Bouras, G., Grigson, S. R., Mirdita, M., Heinzinger, M., Papudeshi, B., Mallawaarachchi, V., Green, R., Kim, R. S., Mihalia, V., Psaltis, A. J., Wormald, P.-J., Vreugde, S., Steinegger, M., & Edwards, R. A. (2026). Protein structure-informed bacteriophage genome annotation with Phold. Nucleic Acids Research, 54(1), gkaf1448. 10.1093/nar/gkaf1448

30. Roux, S., Camargo, A. P., Coutinho, F. H., Dabdoub, S. M., Dutilh, B. E., Nayfach, S., & Tritt, A. (2023). iPHoP: An integrated machine learning framework to maximize host prediction for metagenome-derived viruses of archaea and bacteria. PLOS Biology, 21(4), e3002083. 10.1371/journal.pbio.3002083

31. Chen, G., Jiang, J., & Sun, Y. (2024). RNAVirHost: a machine learning–based method for predicting hosts of RNA viruses through viral genomes. GigaScience, 13, giae059. 10.1093/gigascience/giae059

32. Hulo, C., Castro, E. de, Masson, P., Bougueleret, L., Bairoch, A., Xenarios, I., & Mercier, P. L. (2011). ViralZone: a knowledge resource to understand virus diversity. Nucleic Acids Research, 39(suppl_1), D576–D582. 10.1093/nar/gkq901

33. Duvaud, S., Gabella, C., Lisacek, F., Stockinger, H., Ioannidis, V., & Durinx, C. (2021). Expasy, the Swiss Bioinformatics Resource Portal, as designed by its users. Nucleic Acids Research, 49(W1), W216–W227. 10.1093/nar/gkab225

34. Sayers, E. W., Bolton, E. E., Brister, J. R., Canese, K., Chan, J., Comeau, D. C., Connor, R., Funk, K., Kelly, C., Kim, S., Madej, T., Marchler-Bauer, A., Lanczycki, C., Lathrop, S., Lu, Z., Thibaud-Nissen, F., Murphy, T., Phan, L., Skripchenko, Y., Sherry, S. T. (2021). Database resources of the national center for biotechnology information. Nucleic Acids Research, 50(D1), D20–D26. 10.1093/nar/gkab1112

35. Blum, M., Andreeva, A., Florentino, L. C., Chuguransky, S. R., Grego, T., Hobbs, E., Pinto, B. L., Orr, A., Paysan-Lafosse, T., Ponamareva, I., Salazar, G. A., Bordin, N., Bork, P., Bridge, A., Colwell, L., Gough, J., Haft, D. H., Letunic, I., Llinares-López, F., Bateman, A. (2024). InterPro: the protein sequence classification resource in 2025. Nucleic Acids Research, 53(D1), D444–D456. 10.1093/nar/gkae1082

36. Katoh, K., & Standley, D. M. (2013). MAFFT Multiple Sequence Alignment Software Version 7: Improvements in Performance and Usability. Molecular Biology and Evolution, 30(4), 772–780. 10.1093/molbev/mst010

37. Capella-Gutiérrez, S., Silla-Martínez, J. M., & Gabaldón, T. (2009). trimAl: a tool for automated alignment trimming in large-scale phylogenetic analyses. Bioinformatics, 25(15), 1972–1973. 10.1093/bioinformatics/btp348

38. Minh, B. Q., Schmidt, H. A., Chernomor, O., Schrempf, D., Woodhams, M. D., Haeseler, A. von, & Lanfear, R. (2020). IQ-TREE 2: New Models and Efficient Methods for Phylogenetic Inference in the Genomic Era. Molecular Biology and Evolution, 37(5), 1530–1534. 10.1093/molbev/msaa015

39. Kalyaanamoorthy, S., Minh, B. Q., Wong, T. K. F., Haeseler, A. von, & Jermiin, L. S. (2017). ModelFinder: fast model selection for accurate phylogenetic estimates. Nature Methods, 14(6), 587–589. 10.1038/nmeth.4285

40. Hoang, D. T., Chernomor, O., Haeseler, A. von, Minh, B. Q., & Vinh, L. S. (2018). UFBoot2: Improving the Ultrafast Bootstrap Approximation. Molecular Biology and Evolution, 35(2), 518–522. 10.1093/molbev/msx281

41. Letunic, I., & Bork, P. (2024). Interactive Tree of Life (iTOL) v6: recent updates to the phylogenetic tree display and annotation tool. Nucleic Acids Research, 52(W1), W78–W82. 10.1093/nar/gkae268

42. Sullivan, M. J., Petty, N. K., & Beatson, S. A. (2011). Easyfig: a genome comparison visualizer. Bioinformatics, 27(7), 1009–1010. 10.1093/bioinformatics/btr039

43. Darzentas, N. (2010). Circoletto: visualizing sequence similarity with Circos. Bioinformatics, 26(20), 2620–2621. 10.1093/bioinformatics/btq484

44. Srinivas, P., Peterson, S. B., Gallagher, L. A., Wang, Y., & Mougous, J. D. (2024). Beyond genomics in Patescibacteria: A trove of unexplored biology packed into ultrasmall bacteria. Proceedings of the National Academy of Sciences, 121(51), e2419369121. 10.1073/pnas.2419369121

45. Dion, M. B., Oechslin, F., & Moineau, S. (2020). Phage diversity, genomics and phylogeny. Nature Reviews Microbiology, 18(3), 125–138. 10.1038/s41579-019-0311-5

46. Le, S. Q., & Gascuel, O. (2008). An Improved General Amino Acid Replacement Matrix. Molecular Biology and Evolution, 25(7), 1307–1320. 10.1093/molbev/msn067

47. Zell, R., Groth, M., Selinka, L., & Selinka, H. C. (2024). Diversity of Picorna-Like Viruses in the Teltow Canal, Berlin, Germany. Viruses, 16(7), 1020. 10.3390/v16071020

48. Richard, J. C., Lane, T. W., Agbalog, R. E., Colletti, S. L., Leach, T. C., Dunn, C. D., Bollig, N., Plate, A. R., Munoz, J. T., Leis, E. M., Knowles, S., Standish, I. F., Waller, D. L., & Goldberg, T. L. (2024). Freshwater Mussel Viromes Increase Rapidly in Diversity and Abundance When Hosts Are Released from Captivity into the Wild. Animals, 14(17), 2531. 10.3390/ani14172531

49. Chen, Y. M., Sadiq, S., Tian, J. H., Chen, X., Lin, X. D., Shen, J. J., Chen, H., Hao, Z. Y., Wille, M., Zhou, Z. C., Wu, J., Li, F., Wang, H. W., Yang, W. D., Xu, Q. Y., Wang, W., Gao, W. H., Holmes, E. C., & Zhang, Y. Z. (2022). RNA viromes from terrestrial sites across China expand environmental viral diversity. Nature Microbiology, 7(8), 1312–1323. 10.1038/s41564-022-01180-2

50. Shi, M., Lin, X. D., Tian, J. H., Chen, L. J., Chen, X., Li, C. X., Qin, X. C., Li, J., Cao, J. P., Eden, J. S., Buchmann, J., Wang, W., Xu, J., Holmes, E. C., & Zhang, Y.-Z. (2016). Redefining the invertebrate RNA virosphere. Nature, 540(7634), 539–543. 10.1038/nature20167

51. Elbehery, A. H. A., & Deng, L. (2022). Insights into the global freshwater virome. Frontiers in Microbiology, 13, 953500. 10.3389/fmicb.2022.953500

52. Shade, A., Read, J. S., Youngblut, N. D., Fierer, N., Knight, R., Kratz, T. K., Lottig, N. R., Roden, E. E., Stanley, E. H., Stombaugh, J., Whitaker, R. J., Wu, C. H., & McMahon, K. D. (2012). Lake microbial communities are resilient after a whole-ecosystem disturbance. The ISME Journal, 6(12), 2153–2167. 10.1038/ismej.2012.56

53. Lachnit, T., Thomas, T., & Steinberg, P. (2016). Expanding our Understanding of the Seaweed Holobiont: RNA Viruses of the Red Alga Delisea pulchra. Frontiers in Microbiology, 6, 1489. 10.3389/fmicb.2015.01489

54. Koonin, E. V., Dolja, V. V., Krupovic, M., & Kuhn, J. H. (2021). Viruses Defined by the Position of the Virosphere within the Replicator Space. Microbiology and Molecular Biology Reviews, 85(4), e00193–20. 10.1128/mmbr.00193-20

55. Neri, U., Wolf, Y. I., Roux, S., Camargo, A. P., Lee, B., Kazlauskas, D., Chen, I. M., Ivanova, N., Allen, L. Z., Paez-Espino, D., Bryant, D. A., Bhaya, D., Consortium, R. V. D., Narrowe, A. B., Probst, A. J., Sczyrba, A., Kohler, A., Séguin, A., Shade, A., Gophna, U. (2022). Expansion of the global RNA virome reveals diverse clades of bacteriophages. Cell, 185(21), 4023–4037.e18. 10.1016/j.cell.2022.08.023

56. Edgar, R. C., Taylor, B., Lin, V., Altman, T., Barbera, P., Meleshko, D., Lohr, D., Novakovsky, G., Buchfink, B., Al-Shayeb, B., Banfield, J. F., Peña, M. de la, Korobeynikov, A., Chikhi, R., & Babaian, A. (2022). Petabase-scale sequence alignment catalyses viral discovery. Nature, 602(7895), 142–147. 10.1038/s41586-021-04332-2

57. Tang, X., Shang, J., Chen, G., Chan, K. H. K., Shi, M., & Sun, Y. (2024). SegVir: Reconstruction of Complete Segmented RNA Viral Genomes from Metatranscriptomes. Molecular Biology and Evolution, 41(8), msae171. 10.1093/molbev/msae171

58. Maumus, F., Allen, A. E., Mhiri, C., Hu, H., Jabbari, K., Vardi, A., Grandbastien, M.-A., & Bowler, C. (2009). Potential impact of stress activated retrotransposons on genome evolution in a marine diatom. BMC Genomics, 10(1), 624. 10.1186/1471-2164-10-624

59. Kuroda, K., Yamamoto, K., Nakai, R., Hirakata, Y., Kubota, K., Nobu, M. K., & Narihiro, T. (2022). Symbiosis between Candidatus Patescibacteria and Archaea Discovered in Wastewater-Treating Bioreactors. mBio, 13(5), e01711–22. 10.1128/mbio.01711-22

60. Schulz, F., Yan, Y., Weiner, A. K. M., Ahsan, R., Katz, L. A., & Woyke, T. (2025). Single-cell genomics reveals complex microbial and viral associations in ciliates and testate amoebae. Nature Communications, 16(1), 10336. 10.1038/s41467-025-65263-4

61. Bouderka, F., López-García, P., Deschamps, P., Zhou, Y., Krupovic, M., Gutiérrez-Preciado, A., Ciobanu, M., Bertolino, P., David, G., Moreira, D., & Jardillier, L. (2026). Parasitic connections: a patescibacterial epibiont, its methylotrophic gammaproteobacterial host, and their phages. mBio, 17(4), e00025–26. 10.1128/mbio.00025-26

62. Kumar, A., Nahar, N., Chouhan, D., McLean, J. S., Bor, B., Dong, P. T., & He, X. (2026). A host-encoded prophage targets a Candidate Phyla Radiation bacterium and shapes episymbiotic interactions. Proceedings of the National Academy of Sciences, 123(37), e2615890123. 10.1073/pnas.2615890123

63. Liu, J., Jaffe, A. L., Chen, L., Bor, B., & Banfield, J. F. (2023). Host translation machinery is not a barrier to phages that interact with both CPR and non-CPR bacteria. mBio, 14(6), e01766–23. 10.1128/mbio.01766-23

64. Hu, B., An, L., Wu, M., Xu, J., Nie, Y., & Wu, X. L. (2025). Metagenomics reveals potential interactions between Patescibacteriota and their phages in groundwater ecosystems. mSystems, 11(2), e01204–25. 10.1128/msystems.01204-25

65. Vlok, M., Lang, A. S., & Suttle, C. A. (2019). Marine RNA Virus Quasispecies Are Distributed throughout the Oceans. mSphere, 4(2), 10.1128/mspheredirect.00157-19. 10.1128/mspheredirect.00157-19

